# Maternal transmission gives way to social transmission during gut microbiota assembly in wild mice

**DOI:** 10.1101/2022.08.09.503290

**Authors:** Klara M Wanelik, Aura Raulo, Tanya Troitsky, Arild Husby, Sarah CL Knowles

**Affiliations:** Department of Biology, University of Oxford, Oxford, UK; Department of Computing, University of Turku, Turku, Finland; Department of Ecology and Genetics, Evolutionary Biology Centre, Uppsala University, Sweden

## Abstract

The mammalian gut microbiota influences a wide array of phenotypes and is considered a key determinant of fitness, yet knowledge about the transmission routes by which gut microbes colonise hosts in natural populations remains limited. Here, we use an intensively studied wild population of wood mice (*Apodemus sylvaticus*) to examine how vertical (maternal) and horizontal (social) transmission routes influence gut microbiota composition throughout life. We identify independent signals of maternal transmission (sharing of taxa between a mother and her offspring) and social transmission (sharing of taxa predicted by the social network), whose relative magnitudes shift as hosts age. In early life, gut microbiota composition is predicted to a similar extent by both maternal and social relationships, but by adulthood the impact of maternal transmission becomes undetectable, leaving only a signal of social transmission. By exploring which taxa drive the maternal transmission signal, we further identify a candidate maternally-transmitted bacterial family in wood mice, the *Lactobacillaceae*. Overall, our findings suggest a shifting transmission landscape for wild mice, with a mother’s influence on microbiota composition waning as offspring age, while the impact of social contacts remains strong and consistent.

## Introduction

The gut microbiota has important effects on host phenotypes, from immune development [1], to pathogen resistance [2] and behaviour [3]. An individual’s microbiota is shaped by a variety of forces, including its own genetics [4,5] and diet [6,7], but also the transmission (and subsequent colonisation) of symbiotic gut microbes. Transmission of symbiotic gut microbes can occur at various stages throughout an animal’s life, and from different sources (including the environment, and other conspecifics).

In many species, the mother constitutes an important initial source of microbes that can colonise the gut before, during or after birth. In mammals, although the existence of *in utero* gut microbe transmission is still debated [8], live birth and maternal care provide ample opportunity for vertical transmission during and after birth. Maternal transmission of gut microbes has been well described in humans and inbred laboratory mice, and microbes from the mother’s vaginal, oral, skin, milk and gut microbiota can all be transferred to offspring [9–11]. The vagina constitutes a key source of microbes that can colonise infants during birth [11–14]. In neonates of both mice and humans, the gut microbiota most closely resembles the maternal vaginal microbiota [11,12,15]. The maternal gut is also a key source of gut colonists for young mammals. Well-resolved strain-tracking data from humans indicates that vaginally-derived bacterial strains are transient colonisers and that, among body sites, the maternal gut is the source of the majority of strains transmitted from mother to infant in the first four months of life, including more persistent colonisers of the infant gut [9]. Maternal gut microbes may be transferred through exposure to faeces during birth, through breast milk (as microbes found in breast milk likely originate from the gut [16,17]), via coprophagy [18], or indirectly through nesting material [19]. Although these pathways have been relatively well-studied in humans and laboratory mice, far less is understood about the relative significance, pathways and timing of maternal transmission in wild animals.

Social transmission of microbes from other conspecifics constitutes another important source of transmission that can shape the gut microbiota [20]. Microbes can be shared via direct social interactions such as grooming, aggression or mating, through coprophagy, or via indirect transmission through shared use of space, such as use of the same nest. A growing body of evidence now illustrates the significant influence social transmission can have on the mammalian gut microbiota. For example, in primates, mice and horses, research shows that individuals from the same social group frequently share more microbial taxa and, within social groups or populations, the strength of social interactions predicts microbiota similarity [21–24].

These two major routes of acquiring gut microbes from conspecifics – maternal and social – may each be expected to vary in importance as young animals mature and age. While maternal transmission should be strongest when offspring are interacting closely with their mother (typically in early life), social transmission can take place whenever social interactions are occuring, which for many species constitutes a longer part of the life-course. In mammals, one may expect that social transmission starts later in development and increases as they mature and engage in more, and perhaps different forms of social interaction. However, currently there is only limited data from natural systems to explore this. One human study found no evidence that the impact of social microbial transmission changed with age [25], but studies in wild animals dissecting the relative influence of maternal and social transmission, and how these change with age, are currently lacking.

Accurately detecting and isolating distinct influences of maternal and social transmission on the gut microbiome in free-living animals is challenging. Maternal transmission, for example, is commonly detected by comparing the bacterial taxa present in a mother and her offspring, with a signal of such transmission occurring when mother-offspring pairs share more bacterial taxa than otherwise comparable but genetically unrelated pairs. However, a mother and her offspring may share bacterial taxa for various reasons besides maternal transmission. These include acquisition through exposure to the same environmental reservoir or social contact, or similar (within-host) selection processes that retain or promote the same bacteria, for example through shared genetics or a similar diet. Thus in order to isolate a true signal of maternal transmission, these other confounding variables need to be accounted for. Studies attempting to isolate the impact of maternal transmission in wild systems, where environmental and social influences can freely operate, are rare (but see [26–28]), and among existing studies, the maternal transmission signal is often not robustly isolated from other confounding variables. A second challenge is that the influence of maternal transmission may vary with offspring age. This has been shown in humans and laboratory mice, where the signal of maternal transmission is strongest after birth and gradually decreases over time [29,30]. In wild systems, a signal of maternal transmission could change with offspring age for several non-mutually-exclusive reasons. First, the type, frequency or duration of interaction between a mother and her offspring may change over time, leading to a change in the level of microbial transmission between mother and offspring. For example, a mother may nurse offspring less frequently as they approach weaning, providing less opportunity for microbial transmission. Second, microbial taxa that are maternally transmitted early in life may be outcompeted and replaced by those that are acquired later in life from other sources, for example via social transmission or from the environment. Finally, the infant gut microbiota may go through a process of succession that begins with maternal pioneer microbes, but then may become dominated by other ecological processes such as microbial selection through diet, microbe-microbe interactions and host immunity, making it more difficult to detect the signal of early life maternal transmission.

Here, we use a wild population of wood mice (*Apodemus sylvaticus*) to disentangle the distinct influences of maternal and social transmission on the gut microbiota, and assess how their relative influences change throughout life. We build on our previous work in which we detected a clear signal of social transmission in this population (sharing of taxa being predicted by the social network [22]). Wood mice are well-suited for disentangling the influences of maternal and social transmission, as they are a non-group-living species in which offspring become independent from their mothers after weaning when they emerge from their nest and disperse [31,32] such that social networks are independent of genetic relatedness [22], unlike in many group-living mammals. We use a longitudinal set of faecal samples from a wild population of wood mice for which we also have a rich set of metadata on other covariates that could influence mother-offspring microbiota similarity, including social relationships, spatial locations and seasonality. Using a Bayesian dyadic mixed effects modelling approach, we control for these potential confounders to isolate the effect of maternal transmission and examine how its strength, relative to that of social transmission, changes with age.

## Methods

Data were collected from November 2014 to December 2015 from a wild population of wood mice in a 2.47 ha mixed woodland plot (Nash’s Copse) at Imperial College’s Silwood Park campus, UK. Field methods are described in detail in [22]. Briefly, sterilised live traps were set for one night every 2–4 weeks in an alternating checkerboard design, to ensure even coverage. All traps contained bedding and a standardised bait of 8 peanuts and a slice of apple. At first capture, all mice were injected subcutaneously with a passive integrated transponder tag (PIT-tag) for permanent identification, and a small ear snip was taken for genotyping. At each trapping, demographic data on captured animals was recorded (e.g. sex and age), and faecal samples were collected from traps for gut microbiota analysis and stored within 8 hours of collection at −80°C.

### Social network and space use

Data on mouse space use and the social network was collected in parallel to trapping using a set of nine custom-built PIT-tag loggers distributed across the trapping grid (above ground), as described in [22]. Individuals were considered socially associated with each other if they were detected at the same location on the same night (12 hour period, 6pm to 6am). These data were used to calculate an adjusted version of the Simple Ratio Index (“Adjusted SRI”, see Supplementary information in [22]), that accounts for variable overlap in individual lifespans (i.e. time between first and last logger observation).

### Aging

Individuals were aged as either juvenile, subadult or adult based on pelage and weight (body mass range: juvenile = 10.4 – 14.5g; subadult = 13.0 – 21.5g; adult = 15.1 – 40.4g). Most samples were from adults (*n* = 152 samples) and as we had limited samples from juveniles (*n* = 7 samples) we grouped these with sub-adult samples (*n* = 65 samples) to create one immature age class (*n* = 72 samples).

### Kinship analysis

To derive estimates of host genetic relatedness, ear tissue samples were used to genotype mice at eleven microsatellite loci. A pedigree was then reconstructed using COLONY 2.0.6.5 [33], a program for parental and sibship inference from genotype data. The resulting pedigree was checked against trapping data to remove impossible relationships based on age and trapping date. Finally, kinship results were transformed into genetic relatedness values (unrelated = 0; parent-offspring pair = 0.5; sibling = 0.5; half-sibling = 0.25). Full details of genotyping methods including the target regions chosen and pedigree reconstruction methods are provided in [22].

### Gut microbiota characterisation

The gut microbiota was characterised for 224 faecal samples belonging to 70 genotyped wood mice, including 22 samples from 6 mothers and 37 samples from 17 of their offspring. Microbiota methods for this dataset have been described previously in [22]. Briefly, microbiota profiling involved amplicon sequencing of the 16S rRNA gene (V4 region). Sequence data were processed through the DADA2 pipeline v1.6.0 [34], to infer amplicon sequence variants (ASVs) and taxonomy assigned using the GreenGenes Database (Consortium 13.8). Using the package *phyloseq* [35], ASV counts were normalised to proportional abundance within each sample [36] and singleton ASVs were removed as well as those belonging to non-gut microbial taxa (Cyanobacteria, Mitochondria).

### Statistical analyses

All analyses were conducted in R version 4.0.3 [37]. From the samples characterised, we obtained 4,167 pairs of individuals (of which 15 were mother-offspring pairs), and 24,765 (between-individual) sample-pairs (of which 182 were mother-offspring sample-pairs). To describe compositional microbiota variation, package *vegan* [38] was used to calculate Jaccard distances and Bray-Curtis dissimilarities among samples. We used the Jaccard Index (1-Jaccard distance, the proportion of ASVs shared between a pair of samples) as our primary measure of microbiota similarity, as we considered this metric most relevant for investigating microbial transmission among hosts. However, we also tested whether results were robust to the type of distance metric used, by repeating key analyses using Bray-Curtis similarity (1-Bray-Curtis dissimilarity, which measures similarities in ASV relative abundance as well as presence absence; presented in the Supplementary information).

### Associations between mother-offspring status and microbiota similarity

We used Bayesian (dyadic) regression models in package *brms* [39,40] to model the impact of multiple predictors on pairwise microbiota similarity (Jaccard Index) with a Beta family and logit link. All types of kinship pair were included in our analysis (unrelated pairs, father-offspring pairs, mother-offspring pairs, sibling pairs and half-sibling pairs) and the binary predictor mother-offspring status (that captured whether a pair of individuals were a mother-offspring pair, 1, or not, 0), tested for an effect of maternal transmission. Since individual mice were often sampled multiple times, and all individuals were included in multiple pairwise similarity measures, we included two multi-membership random effects to account for the non-independence inherent to this type of data: (1) a random intercept term for the individuals in each dyad (Individual A + Individual B) and (2) a random intercept term for the samples in each dyad (Sample A + Sample B). As in [22], we included other variables that were either suspected or previously demonstrated drivers of dyadic microbiota similarity. Since we used a dyadic framework, these predictor variables were coded as similarities, distances or associations, describing a *pair* of samples. These included genetic relatedness (unrelated pair = 0; parent-offspring pair = 0.5; sibling pair = 0.5; half-sibling pair = 0.25), social association (adjusted SRI), age class similarity (same age class = 1; different age class = 0), sex similarity (same sex = 1; different sex = 0), spatial distance (distance between individuals’ mean spatial coordinates from logger records) and sampling interval (time in days between which samples were taken). We excluded all pairs of samples where both individuals were immature (immature-immature pairs; *n* = 2,566 sample-pairs), leaving only adult-immature and adult-adult pairs. To allow us to test whether the mother-offspring effect (signal of maternal transmission) varied with offspring age class, we also excluded a small number of mother-offspring sample-pairs where the mother was sampled as an immature (*n* = 16 sample-pairs). This meant that age class similarity among mother-offspring pairs was a function of offspring age class only i.e. mother-offspring pairs could only differ in age class (adult-immature pair) if offspring were immature and share the same age class (adult-adult pair) if both mother and offspring were adult. We then included an interaction term between the variable mother-offspring status and age class similarity in order to test whether the maternal transmission effect varied with offspring age. The final number of sample-pairs included in the analysis was 22,199 sample-pairs (of which 170 were mother-offspring sample-pairs). All models were checked for convergence by visual inspection of trace plots and the *Rhat* statistic.

### Identifying which bacterial taxa associate with mother-offspring status

We used the same approach used to identify candidate socially-transmitted bacterial taxa in [22], to here identify candidate maternally transmitted bacterial taxa. We tested how each bacterial family influenced the effect size of (1) the main effect of mother-offspring status, and (2) the interaction between mother-offspring status and age class similarity. We recalculated the Jaccard Index excluding each bacterial family in turn, then compared effect sizes and credible intervals from the same models run in the package *MCMCglmm* [41] using these indices.

## Results

### Identifying maternal and social transmission signals

We found no association between mother-offspring status and social association (Mantel test: *r* = 0.01, *p* = 0.17), allowing us to dissect the distinct influences of maternal and social transmission on the gut microbiota. Across the whole dataset, we identified a clear signal of maternal transmission in wild mice, since mother-offspring status positively predicted the proportion of shared gut microbial ASVs (posterior mean = 0.21, 95% CI = 0.13, 0.30; Table S1), when controlling for a range of confounding variables that could also influence mother-offspring microbiota similarity (social association, spatial distance, sampling interval and genetic relatedness). This maternal transmission effect was weaker overall than the social transmission effect (posterior mean = 0.32, 95% CI = 0.24, 0.41; Table S1). Consistent with [22], we found that other variables also predicted microbiota similarity, including spatial distance and sampling interval (Table S1). Additionally, we found that genetic relatedness predicted microbiota similarity (unlike [22], where an absence of such a relationship was reported). Contrary to expectation, we found that among non-mother-offspring pairs, more closely related individuals had less similar microbiota (Table S1). This is unlikely to be a genetic effect, and more likely to be caused by some environmental dissimilarity between other (related but not mother-offspring) kinship pairs and requires further investigation. Consistent results were obtained when using Bray-Curtis similarities (Table S2).

### The influence of age class on maternal and social transmission signals

When an interaction term between mother-offspring status and age class similarity was included in the model, this interaction term was significant (posterior mean = −0.27, 95% CI = −0.41, −0.13; Table S1), indicating that the mother-offspring effect was stronger when offspring were immature. Among adult-immature pairs, mother-offspring pairs shared a higher proportion of ASVs than non-mother-offspring pairs (posterior mean for mother-offspring status = 0.36, 95% CI = 0.24, 0.47; Table S1), whereas among adult-adult pairs, mother-offspring and non-mother-offspring pairs shared a similar proportion of ASVs (posterior mean for mother-offspring status = 0.09, 95% CI = −0.02, 0.20; Table S1; Fig. 1). We also included an interaction term between social-association strength and age class similarity, but this was not significant suggesting that the signal of social transmission remained consistent throughout life (posterior mean = 0.07, 95% CI = −0.08, 0.22; Table S1). Although we found that the maternal transmission effect was weaker than the social transmission effect overall (see above), when separating by age class, social and maternal transmission had equally strong effects on microbiota composition early in life, with the relative influence of maternal transmission waning in adulthood and leaving only a signal of social transmission (Fig. 2). Again, consistent results were obtained when using Bray-Curtis similarities (Table S2).

**Figure 1.**
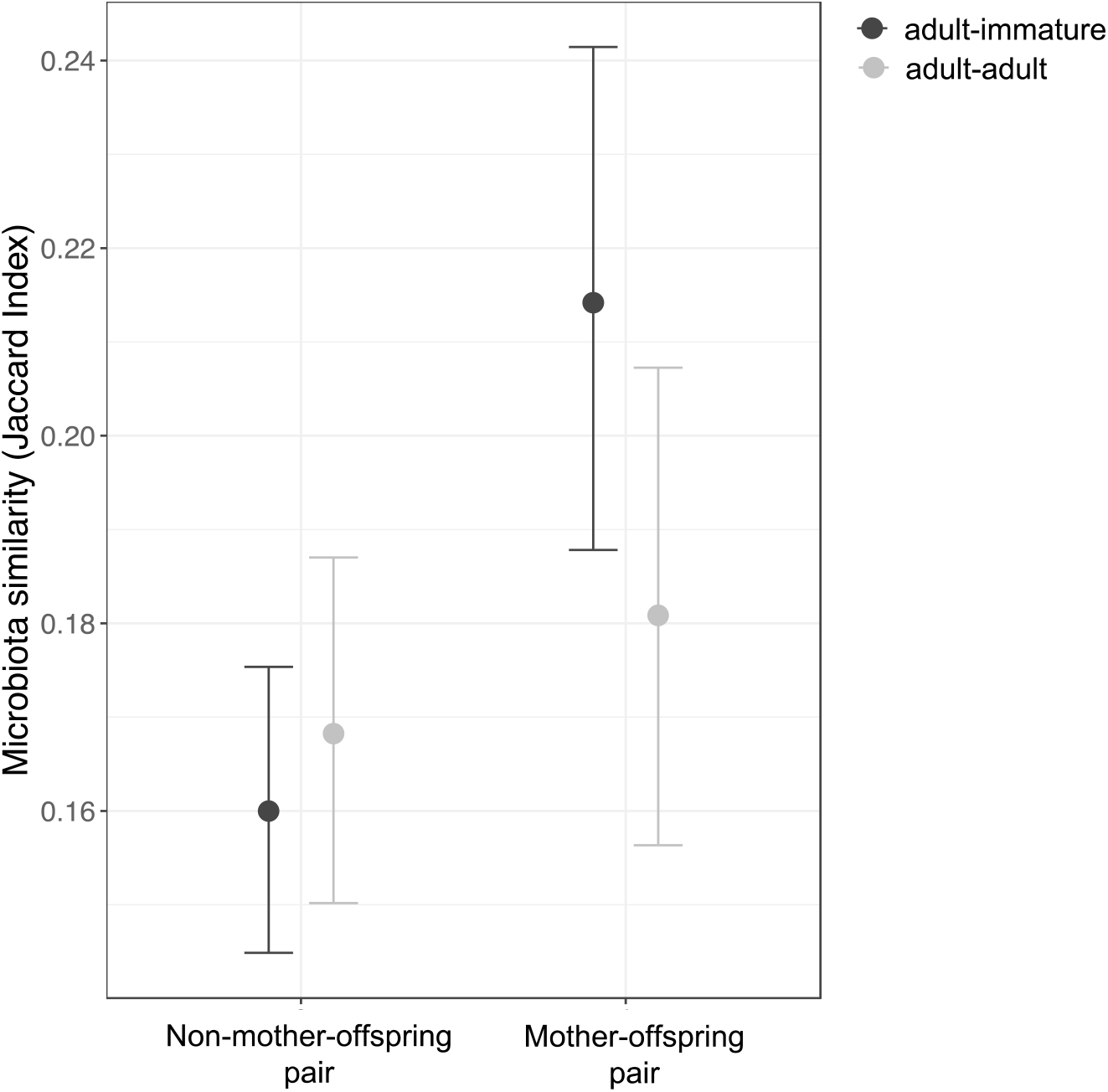
The maternal transmission signal depends on age class similarity (adult-immature or adult-adult pair). Estimates (points) and their 95% credible intervals are plotted from Bayesian regression (*brms*) models controlling for other confounding variables, with pairwise microbiota similarity among hosts (Jaccard Index) as the response. Among mother-offspring pairs where one is an adult and the other immature (adult-immature pairs), the immature individual is always the offspring. Non-mother-offspring pairs include all other types of kinship pairs (unrelated pairs, father-offspring pairs, sibling pairs and half-sibling pairs).

**Figure 2.**
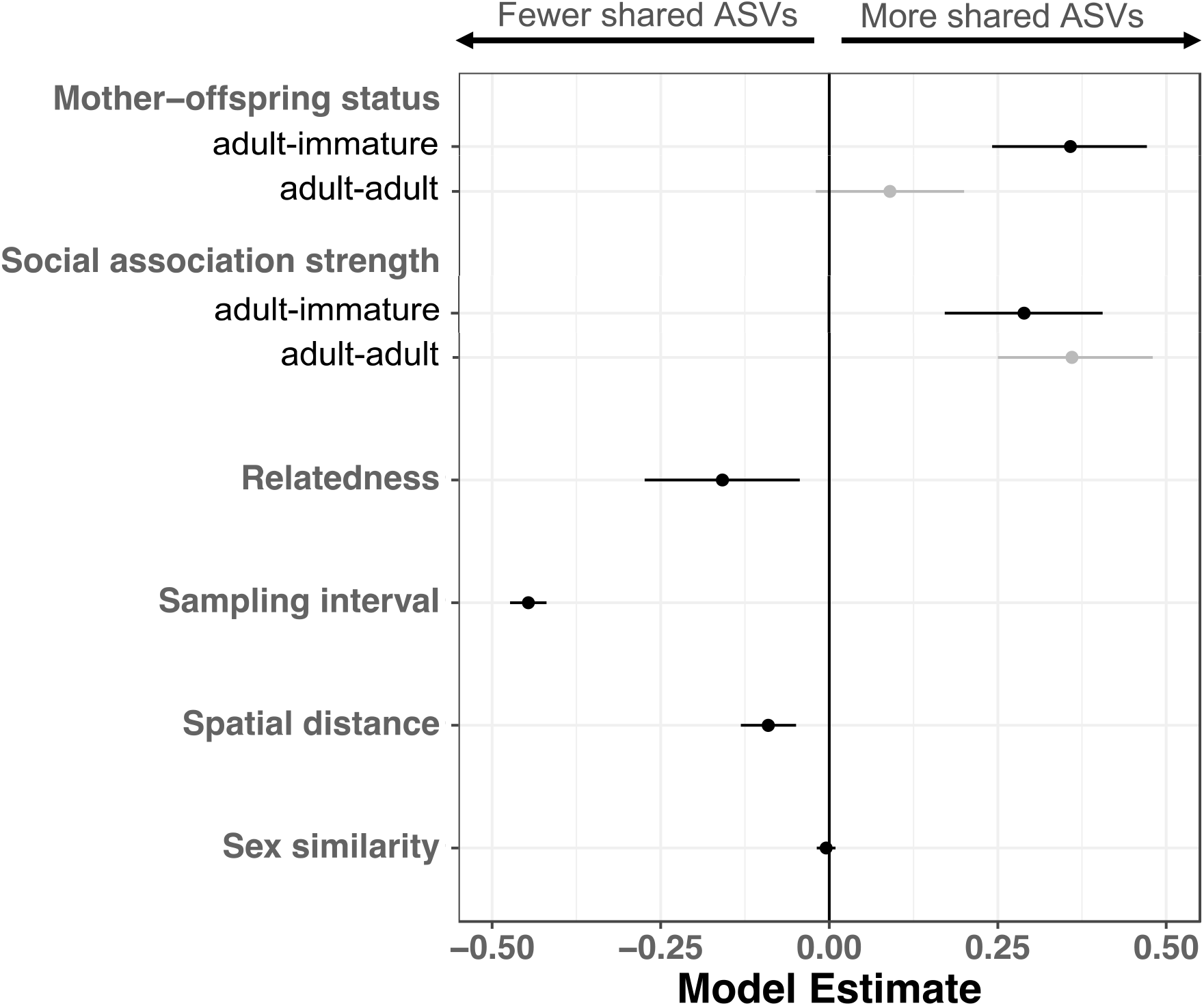
Maternal and social transmission predict gut microbiota similarity equally strong early in life (adult-immature pair), with the relative influence of maternal transmission waning in adulthood (adult-adult pair). Social association strength was estimated using an adjusted version of the Simple Ratio Index (“Adjusted SRI”) using a 12-hour, overnight time window. Effect size estimates (points) and their 95% credible intervals are plotted from Bayesian regression (*brms*) models with pairwise microbiota similarity among hosts (Jaccard Index) as the response. Where credible intervals do not overlap zero, a variable significantly predicts microbiota similarity.

### Identifying candidate maternally-transmitted bacterial taxa

Both the main effect of mother-offspring status and the interaction effect between mother-offspring status and age class similarity remained significant or marginally significant in all models where a single bacterial family was excluded, suggesting that they did not depend entirely on any single bacterial family (Fig. 3). However, excluding the *Lactobacillaceae* did weaken both effect sizes considerably (main effect: taking the posterior mean from 0.03 to 0.01; interaction effect: taking the posterior mean from - 0.04 to −0.02; Fig. 3).

**Figure 3.**
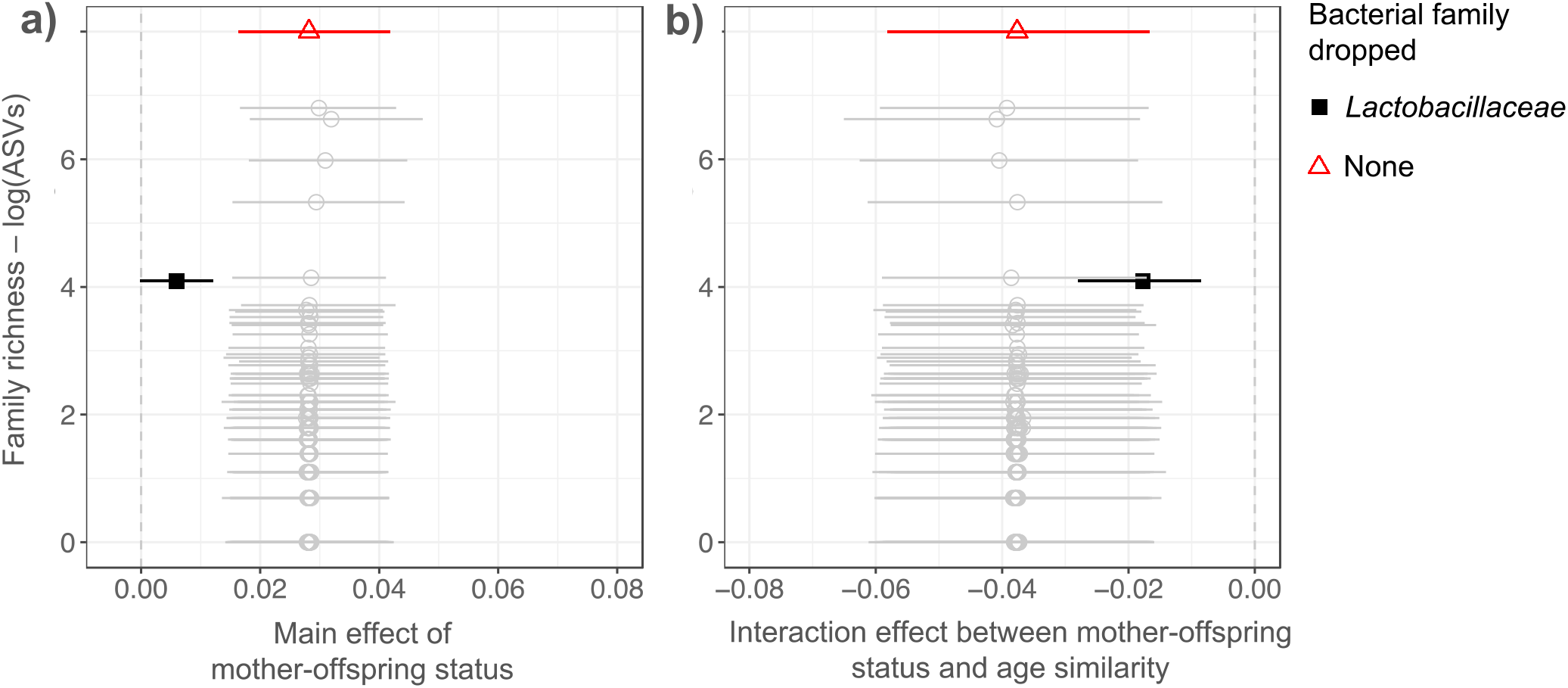
The influence of specific bacterial families on the maternal transmission signal. Effect sizes for (a) the main effect of mother-offspring status, and (b) the interaction between mother-offspring status and age class similarity and 95% credible intervals are plotted from 146 Bayesian regression models, in each of which a single bacterial family was excluded. Effects are plotted against the species richness of each dropped family (logged number of ASVs). Effect sizes and their 95% credible intervals from the full model (with no bacterial families dropped) are included for reference at the top.

## Discussion

In this study, we examine the relative influence of maternal and social transmission on the gut microbiota as hosts age in a wild mouse population. We do this by drawing on a rare example of a longitudinal dataset of paired mother-offspring faecal samples taken from the wild, for which we also have a rich set of metadata. By controlling for other confounding variables we are able to isolate distinct maternal and social transmission signals. We show that who is important in shaping an offspring’s gut microbiota composition changes through life – while the importance of the mother declines, the importance of other social contacts remains stable from the early independent phase into adulthood. In doing so, we also describe, for the first time, maternal transmission of gut microbiota in wild mice, suggesting that the maternal transmission signal observed in laboratory (inbred) mice [10,11] extends to their wild counterparts, and remains detectable post-weaning. Our estimate of the maternal transmission signal is likely conservative as all the offspring we sampled as part of this study were weaned (and trappable). Despite this, we still found a strong maternal transmission signal, which may be even stronger at or shortly after birth, had we been able to sample at this point. Similarly, we cannot say from our data how much influence social transmission might have at the very earliest stage of life (pre-weaning), but expect this to be minimal since before weaning, wood mice are raised solely by their mother in underground burrows, and mothers are territorial during this time [42]. What we can say, is that by the time individuals are weaned, independent and capable of being caught in live traps, social transmission already has an important influence.

Of course, some forms of maternal transmission may be considered somewhat ‘social’ in nature, driven by close proximity between a mother and her offspring within the nest. A close link between maternal and social transmission is consistent with our finding that the same bacterial family, the *Lactobacillaceae*, has been found here to influence the maternal transmission signal, as well as having previously been shown to influence the social transmission signal [22]. The *Lactobacillaceae* are non-spore forming and anaerobic [43], which means they cannot persist for any length of time in the environment. These life history traits are consistent with this taxon being transmitted between individuals in close proximity, either from a mother to her offspring below ground, or from one conspecific to another above ground. Previous studies have also identified other maternally-transmitted taxa. In humans, it is well established that bifidobacteria are maternally transmitted [44–47]. More recently, bifidobacteria have been shown to be commonly maternally transmitted across a wide range of mammals (including primates and non-primates) [44]. Although bifidobacteria were not identified as important for maternal transmission in this study, they were detected in the wood mouse gut microbiota. Other candidate maternally-transmitted taxa include *Lactobacillus* [48] and *Bacteroides* [11,27]. All these taxa are implicated in the degradation of milk oligosaccharides. Within the *Lactobacillaceae* family, the most common genus within our mouse population was *Lactobacillus*. Gut-associated *Lactobacillus* is found in human breastmilk, can be transmitted to neonates via breastfeeding [17] and is involved in milk breakdown [14]. We might then expect this taxon to be transitional, giving way to other taxa (e.g. fibre fermenters) as diet changes in later life. Indeed, transitional colonisation by *Lactobacillus* has been observed in humans [48]. Further work is needed to explore whether an age-related decline in *Lactobacillus* might explain the overall age-related decline in strength of the maternal transmission signal we have identified here, and whether specific maternally-transmitted taxa in this family play important functional roles in early life.

Transmission events early in life may have long-lasting effects on an individual. In terms of shared taxa, we found that adult offspring were no more similar to their mother than to any other individual in the population. However, maternal impacts on an adult’s microbiota that are harder to detect may nonetheless persist, if maternal transmission events early in life give rise to predictable patterns of microbial succession that permanently change a microbial community. Our analyses would not detect such a change, but higher resolution time-series analyses combined with AI could potentially be used to test this hypothesis in the future. Even transient changes in microbiota early in life can shape development in key ways. For example, a recent study suggested that such transient changes may help establish alternative development trajectories in industrialised and non-industrialised human populations [49]. Short-term antibiotic-driven microbiota perturbation in early life has also been linked to long-term host metabolic effects implicated in obesity [50], and long-term immunological effects implicated in asthma and other autoimmune diseases [51]. For example, the genus *Lactobacillus* (in the family *Lactobacillaceae*, here identified as a candidate maternally-transmitted family) is thought to play a role in the priming and maturation of dendritic cells, which initiate and shape the adaptive immune response [52]. Thus microbiota differences involving this genus early in life could have important knock-on effects on the functioning of the adaptive immune response – a key fitness determinant. Future wild studies that assess links between early-life microbial communities and later life traits associated with health or fitness (similar to [53]) would be useful in this respect. We suggest wild mouse systems, such as that used here, are a tractable system in which to quantify these long-term, and potentially profound, effects of maternal transmission.

## Author contributions

SCLK conceived and designed the field study, carried out fieldwork, collected samples and cleaned field data for analysis. AR conducted microbiome laboratory work. KMW conceived the analyses, analysed the data and led writing of the manuscript. TT and AH performed mouse genotyping and built the pedigree. TT, AR and SCLK provided input during analyses, and all authors contributed to and reviewed the manuscript.

## Acknowledgements

We thank Aurelio Malo for loan of the PIT-tag loggers and advice on field experimental design, Bryony Allen, Saskia Ricks, Terence Chung, Selene Guitierrez Al-Khudhairy, Alice Weightman, Rohan Raval and Joe Williamson for assistance with fieldwork, and Mike Francis for manufacture of loggers and support with their use. We thank Paul Buerkner for help with *brms* models.

## References

1. Sanidad KZ, Zeng MY.s Neonatal gut microbiome and immunity. Current Opinion in Microbiology. 2020;56:30–7.

2. Kamada N, Chen GY, Inohara N, Núñez G. Control of pathogens and pathobionts by the gut microbiota. Nature Immunology. 2013;14:685–90.

3. Davidson GL, Raulo A, Knowles SCL. Identifying Microbiome-Mediated Behaviour in Wild Vertebrates. Trends in Ecology and Evolution. 2020;35:972–80.

4. Bonder MJ, Kurilshikov A, Tigchelaar EF, Mujagic Z, Imhann F, Vila AV, et al. The effect of host genetics on the gut microbiome. Nature Genetics. 2016;48:1407–12.

5. Benson AK, Kelly SA, Legge R, Ma F, Low SJ, Kim J, et al. Individuality in gut microbiota composition is a complex polygenic trait shaped by multiple environmental and host genetic factors. Proc Natl Acad Sci U S A. 2010;107:18933–8.

6. Turnbaugh PJ, Ridaura VK, Faith JJ, Rey FE, Knight R, Gordon JI. The effect of diet on the human gut microbiome: A metagenomic analysis in humanized gnotobiotic mice. Science Translational Medicine. 2009;1:6ra14.

7. David LA, Maurice CF, Carmody RN, Gootenberg DB, Button JE, Wolfe BE, et al. Diet rapidly and reproducibly alters the human gut microbiome. Nature. 2014;505:559–63.

8. Perez-Muñoz ME, Arrieta MC, Ramer-Tait AE, Walter J. A critical assessment of the “sterile womb” and “in utero colonization” hypotheses: Implications for research on the pioneer infant microbiome. Microbiome. 2017;5.

9. Ferretti P, Pasolli E, Tett A, Asnicar F, Gorfer V, Fedi S, et al. Mother-to-Infant Microbial Transmission from Different Body Sites Shapes the Developing Infant Gut Microbiome. Cell Host and Microbe. 2018;24:133–45.

10. Moeller AH, Suzuki TA, Phifer-Rixey M, Nachman MW. Transmission modes of the mammalian gut microbiota. Science. 2018;362:453–7.

11. Koo H, McFarland BC, Hakim JA, Crossman DK, Crowley MR, Rodriguez JM, et al. An individualized mosaic of maternal microbial strains is transmitted to the infant gut microbial community. Royal Society Open Science. 2020;7:192200.

12. Dominguez-Bello MG, Costello EK, Contreras M, Magris M, Hidalgo G, Fierer N, et al. Delivery mode shapes the acquisition and structure of the initial microbiota across multiple body habitats in newborns. Proc Natl Acad Sci U S A. 2010;107:11971–5.

13. Biasucci G, Rubini M, Riboni S, Morelli L, Bessi E, Retetangos C. Mode of delivery affects the bacterial community in the newborn gut. Early Human Development. 2010;86:13–5.

14. Bäckhed F, Roswall J, Peng Y, Feng Q, Jia H, Kovatcheva-Datchary P, et al. Dynamics and stabilization of the human gut microbiome during the first year of life. Cell Host and Microbe. 2015;17:690–703.

15. Pantoja-Feliciano IG, Clemente JC, Costello EK, Perez ME, Blaser MJ, Knight R, et al. Biphasic assembly of the murine intestinal microbiota during early development. ISME Journal. 2013;7:1112–5.

16. Rodríguez JM. The Origin of Human Milk Bacteria: Is There a Bacterial Entero-Mammary Pathway during Late Pregnancy and Lactation? American Society for Nutrition Adv Nutr. 2014;5:779–84.

17. Jost T, Lacroix C, Braegger CP, Rochat F, Chassard C. Vertical mother-neonate transfer of maternal gut bacteria via breastfeeding. Environmental Microbiology. 2014;16:2891–904.

18. Osawa R, Blanshard WH, O’Callaghan PG. Microbiological studies of the intestinal microflora of the koala, Phascolarctos cinereus. II. Pap, a Special Maternal Faeces Consumed by Juvenile Koalas. Australian Journal of Zoology. 1993;41:527–36.

19. Campos-Cerda F, Bohannan BJM. The Nidobiome: A Framework for Understanding Microbiome Assembly in Neonates. Trends in Ecology and Evolution. Elsevier Ltd; 2020;35:573–82.

20. Sarkar A, Harty S, Johnson KVA, Moeller AH, Archie EA, Schell LD, et al. Microbial transmission in animal social networks and the social microbiome. Nature Ecology and Evolution. 2020;4:1020–35.

21. Tung J, Barreiro LB, Burns MB, Grenier JC, Lynch J, Grieneisen LE, et al. Social networks predict gut microbiome composition in wild baboons. Elife. 2015;4:e05224.

22. Raulo A, Allen B, Troitsky T, Husby A, Firth JA, Coulson T, et al. Social networks strongly predict the gut microbiota of wild mice. ISME Journal. 2021;15:2601–13.

23. Dill-McFarland KA, Tang ZZ, Kemis JH, Kerby RL, Chen G, Palloni A, et al. Close social relationships correlate with human gut microbiota composition. Scientific Reports. 2019;9:703.

24. Stothart MR, Greuel RJ, Gavriliuc S, Henry A, Wilson AJ, McLoughlin PD, et al. Bacterial dispersal and drift drive microbiome diversity patterns within a population of feral hindgut fermenters. Molecular Ecology. 2020;30:555–71.

25. Brito IL, Gurry T, Zhao S, Huang K, Young SK, Shea TP, et al. Transmission of human-associated microbiota along family and social networks. Nature Microbiology. 2019;4:964–71.

26. Reese AT, Phillips SR, Owens LA, Venable EM, Langergraber KE, Machanda ZP, et al. Age Patterning in Wild Chimpanzee Gut Microbiota Diversity Reveals Differences from Humans in Early Life. Current Biology. 2021;31:613–20.

27. Baniel A, Petrullo L, Mercer A, Reitsema L, Sams S, Jacinta C, et al. Maternal effects on early-life gut microbiome maturation in a wild nonhuman primate. bioRxiv. 2021.

28. Ren T, Boutin S, Humphries MM, Dantzer B, Gorrell JC, Coltman DW, et al. Seasonal, spatial, and maternal effects on gut microbiome in wild red squirrels. Microbiome; 2017;5:163.

29. Korpela K, Costea P, Coelho LP, Kandels-Lewis S, Willemsen G, Boomsma DI, et al. Selective maternal seeding and environment shape the human gut microbiome. Genome Research. 2018;28:561–8.

30. Friswell MK, Gika H, Stratford IJ, Theodoridis G, Telfer B, Wilson ID, et al. Site and strain-specific variation in gut microbiota profiles and metabolism in experimental mice. PLoS ONE. 2010;5:e8584.

31. Flowerdew JR. Field and Laboratory Experiments on the Social Behaviour and Population Dynamics of the Wood Mouse (Apodemus sylvaticus). The Journal of Animal Ecology. 1974;43:499–511.

32. Watts CHS. The Regulation of Wood Mouse (Apodemus sylvaticus) Numbers in Wytham Woods, Berkshire. The Journal of Animal Ecology. 1969;38:285–304.

33. Wang J, Santure AW. Parentage and sibship inference from multilocus genotype data under polygamy. Genetics. 2009;181:1579–94.

34. Callahan BJ, McMurdie PJ, Rosen MJ, Han AW, Johnson AJA, Holmes SP. DADA2: High-resolution sample inference from Illumina amplicon data. Nature Methods. 2016;13:581–3.

35. McMurdie PJ, Holmes S. Phyloseq: An R Package for Reproducible Interactive Analysis and Graphics of Microbiome Census Data. PLoS ONE. 2013;8:e61217.

36. McKnight DT, Huerlimann R, Bower DS, Schwarzkopf L, Alford RA, Zenger KR. Methods for normalizing microbiome data: An ecological perspective. Methods in Ecology and Evolution. 2019;10:389–400.

37. R Core Team. R: A language and environment for statistical computing. R Foundation for Statistical Computing. Vienna, Austria; 2020. Available from: https://www.r-project.org/

38. Oksanen K, Guillaume Blanchet F, Friendly M, Kindt R, Legendre P, McGlinn D, et al. vegan: Community Ecology Package. R package version 2.5-7. 2020. Available from: https://cran.r-project.org/package=vegan

39. Bürkner PC. Advanced Bayesian multilevel modeling with the R package brms. R Journal. 2018;10:395–411.

40. Bürkner PC. brms: An R package for Bayesian multilevel models using Stan. Journal of Statistical Software. 2017;80.

41. Hadfield JD. MCMCglmm: MCMC Methods for Multi-Response GLMMs in R. Journal of Statistical Software. 2010;33.

42. Wolton R. The ranging and nesting behaviour of Wood mice, Apodemus sylvaticus (Rodentia: Muridae), as revealed by radio-tracking. Journal of Zoology. 1985;206:203–22.

43. Schleifer K. Lactobacillaceae. In: DeVos P, Dedysh S, Hedlund B, Kämpfer P, Rainey F, Trujillo M, et al., editors. Bergey’s Manual of Systematics of Archaea and Bacteria. Hoboken, New Jersey: John Wiley & Sons; 2015.

44. Milani C, Mangifesta M, Mancabelli L, Lugli GA, James K, Duranti S, et al. Unveiling bifidobacterial biogeography across the mammalian branch of the tree of life. ISME Journal. 2017;11:2834–47.

45. Milani C, Mancabelli L, Lugli GA, Duranti S, Turroni F, Ferrario C, et al. Exploring vertical transmission of bifidobacteria from mother to child. Applied and Environmental Microbiology. 2015;81:7078–87.

46. Avershina E, Lundgård K, Sekelja M, Dotterud C, Storrø O, Øien T, et al. Transition from infant-to adult-like gut microbiota. Environ Microbiol. 2016;18:2226–36.

47. Duranti S, Lugli GA, Mancabelli L, Armanini F, Turroni F, James K, et al. Maternal inheritance of bifidobacterial communities and bifidophages in infants through vertical transmission. Microbiome; 2017;5:66.

48. Dotterud CK, Avershina E, Sekelja M, Simpson MR, Rudi K, Storrø O, et al. Does maternal perinatal probiotic supplementation alter the intestinal microbiota of mother and child? Journal of Pediatric Gastroenterology and Nutrition. 2015;61:200–7.

49. Olm MR, Dahan D, Carter MM, Merrill BD, Yu B, Jain S, et al. Robust Variation in Infant Gut Microbiome Assembly Across a Spectrum of Lifestyles. Science. 2022;376:1220–3.

50. Cox LM, Yamanishi S, Sohn J, Alekseyenko A v., Leung JM, Cho I, et al. Altering the intestinal microbiota during a critical developmental window has lasting metabolic consequences. Cell. 2014;158:705–21.

51. Russell SL, Gold MJ, Hartmann M, Willing BP, Thorson L, Wlodarska M, et al. Early life antibiotic-driven changes in microbiota enhance susceptibility to allergic asthma. EMBO Reports. 2012;13:440–7.

52. Smits HH, Engering A, van der Kleij D, de Jong EC, Schipper K, van Capel TMM, et al. Selective probiotic bacteria induce IL-10-producing regulatory T cells in vitro by modulating dendritic cell function through dendritic cell-specific intercellular adhesion molecule 3-grabbing nonintegrin. Journal of Allergy and Clinical Immunology. 2005;115:1260–7.

53. Wanelik KM, Begon M, Bradley JE, Friberg IM, Christopher H. Early-life immune expression profiles predict later life health and fitness in a wild rodent. bioRxiv. 2021.

