## Supplementary information for "Maternal transmission gives way to social transmission during gut microbiota assembly in wild mice"

This additional information accompanies:

**Table S1** Results of *brms* models testing the effect of mother-offspring status and covariates on microbiota similarity (Jaccard Index). Significant terms (where 95% credible intervals do not include zero) are shown in bold. Est. Error indicates the standard deviation of the posterior distribution.

| **Without interaction terms** | | | | |
|---|---|---|---|---|
| | **Estimate** | **Est. Error** | **l-95% CI** | **u-95% CI** |
| Intercept | -1.46 | 0.06 | -1.58 | -1.35 |
| Sex similarity | 0.00 | 0.01 | -0.02 | 0.01 |
| **Spatial distance** | **-0.10** | **0.02** | **-0.14** | **-0.06** |
| **Temporal distance** | **-0.45** | **0.01** | **-0.47** | **-0.42** |
| **Relatedness** | **-0.16** | **0.06** | **-0.28** | **-0.05** |
| **Social association strength** | **0.32** | **0.04** | **0.24** | **0.41** |
| **Mother-offspring status** | **0.21** | **0.04** | **0.13** | **0.30** |
| Age class similarity | 0.06 | 0.05 | -0.04 | 0.15 |
| **With interaction terms** | | | | |
| | **Estimate** | **Est. Error** | **l-95% CI** | **u-95% CI** |
| Intercept | -1.46 | 0.06 | -1.58 | -1.36 |
| Sex similarity | 0.00 | 0.01 | -0.02 | 0.01 |
| **Spatial distance** | **-0.09** | **0.02** | **-0.13** | **-0.05** |
| **Temporal distance** | **-0.45** | **0.01** | **-0.47** | **-0.42** |
| **Relatedness** | **-0.16** | **0.06** | **-0.27** | **-0.04** |
| **Social association strength** | **0.29** | **0.06** | **0.17** | **0.41** |
| Mother-offspring status | 0.36 | 0.06 | 0.24 | 0.47 |
| Age class similarity | 0.06 | 0.05 | -0.04 | 0.15 |
| Social association strength:Age class similarity | 0.07 | 0.08 | -0.08 | 0.22 |

| | | | | |
|---|---|---|---|---|
| **Mother-offspring status:Age class similarity** | **-0.27** | **0.07** | **-0.41** | **-0.13** |

**Table S2** Results of *brms* models testing the effect of mother-offspring status and covariates on microbiota similarity (Bray-Curtis). Significant terms (where 95% credible intervals do not include zero) are shown in bold. Est. Error indicates the standard deviation of the posterior distribution.

| **Without interaction terms** | | | | |
|---|---|---|---|---|
| | **Estimate** | **Est. Error** | **l-95% CI** | **u-95% CI** |
| Intercept | -0.79 | 0.06 | -0.91 | -0.68 |
| Sex similarity | 0.00 | 0.01 | -0.02 | 0.01 |
| **Spatial distance** | **-0.10** | **0.02** | **-0.14** | **-0.06** |
| **Temporal distance** | **-0.46** | **0.01** | **-0.49** | **-0.43** |
| **Relatedness** | **-0.17** | **0.06** | **-0.28** | **-0.06** |
| **Social association strength** | **0.35** | **0.05** | **0.26** | **0.44** |
| **Mother-offspring status** | **0.22** | **0.04** | **0.13** | **0.31** |
| Age class similarity | 0.05 | 0.05 | -0.04 | 0.15 |
| **With interaction terms** | | | | |
| | **Estimate** | **Est. Error** | **l-95% CI** | **u-95% CI** |
| Intercept | -0.79 | 0.06 | -0.91 | -0.68 |
| Sex similarity | 0.00 | 0.01 | -0.02 | 0.01 |
| **Spatial distance** | **-0.09** | **0.02** | **-0.14** | **-0.05** |
| **Temporal distance** | **-0.46** | **0.01** | **-0.49** | **-0.43** |
| **Relatedness** | **-0.17** | **0.06** | **-0.28** | **-0.06** |
| Social association strength | 0.32 | 0.06 | 0.20 | 0.45 |
| Mother-offspring status | 0.36 | 0.06 | 0.25 | 0.48 |

| | | | | |
|---|---|---|---|---|
| Age class similarity | 0.06 | 0.05 | -0.04 | 0.15 |
| Social association strength:Age class similarity | 0.06 | 0.08 | -0.10 | 0.22 |
| **Mother-offspring status:Age class similarity** | **-0.27** | **0.07** | **-0.40** | **-0.13** |
